# Colon-specific immune microenvironment regulates cancer progression versus rejection

**DOI:** 10.1101/2020.01.02.892711

**Authors:** Giulia Trimaglio, Anne-Françoise Tilkin-Mariamé, Virginie Feliu, Françoise Lauzeral-Vizcaino, Marie Tosolini, Carine Valle, Maha Ayyoub, Olivier Neyrolles, Nathalie Vergnolle, Yoann Rombouts, Christel Devaud

## Abstract

**Background:** Immunotherapies have achieved clinical benefit in many types of cancer but remain limited to a subset of patients in colorectal cancer (CRC). Resistance to immunotherapy can be attributed in part to tissue-specific factors constraining antitumor immunity. Thus, a better understanding of how the colon microenvironment shapes the immune response to CRC is needed to identify mechanisms of resistance to immunotherapies and guide the development of novel therapeutics.

**Methods:** In an orthotopic mouse model of MC38-CRC, tumor progression was monitored by bioluminescence imaging and the immune signatures were assessed at a transcriptional level using NanoString and at a cellular level by flow cytometry.

**Results:** Despite initial tumor growth in all mice, only 25 to 35% of mice developed a progressive lethal CRC while the remaining animals spontaneously rejected their solid tumor. No tumor rejection was observed in the absence of adaptive immunity, nor when MC38 cells were injected in non-orthotopic locations, subcutaneously or into the liver. We observed that progressive CRC tumors exhibited a protumor immune response, characterized by a regulatory T-lymphocyte pattern, discernible shortly post-tumor implantation, as well as suppressive myeloid cells. In contrast, tumor-rejecting mice presented an early inflammatory response and an antitumor microenvironment enriched in CD8^+^ T cells.

**Conclusions:** Taken together, our data demonstrate the role of the colon microenvironment in regulating the balance between anti or protumor immune responses. While emphasizing the relevance of the CRC orthotopic model, they set the basis for exploring the impact of the identified signatures in colon cancer response to immunotherapy.

## Introduction

Tumor-infiltrating innate and adaptive immune cells play a dual role in cancer development [1]. In a first phase called “elimination”, immune cells can recognize and kill recently transformed malignant cells. During a second “equilibrium” phase, the rare tumor variants that have survived the elimination can enter a non-growing dormant state that can last for long periods of time [2]. Finally, in a third “escape” phase, tumor cells exit dormancy and proliferate again with the help of the immunosuppressive microenvironment [2]. The antitumor immune response predominantly relies on tumor antigen-specific effector CD8^+^ T lymphocytes and other lymphoid cell subsets, while the protumor axis mainly involves immunosuppressive regulatory T cells (Treg), myeloid-derived suppressor cells (MDSC) and anti-inflammatory type 2 macrophages (M2) [1]. In this context, therapies that harness and enhance antitumor effector cells, such as immune checkpoint blockade therapies, have led to clinical benefit in several malignancies including melanoma, non-small cell lung cancer, and renal cell carcinoma [3].

While CRC remains the third most prevalent cause of cancer-related deaths worldwide [4], the current success of immunotherapy is limited to ~5% of all CRC patients [3]. Patients responding to immunotherapy exhibit a defective DNA mismatch repair system (MMR)/microsatellite Instability-High (MSI) CRC phenotype that may have higher immunological potential [5]. More recently, it has been demonstrated that the type, location and density of adaptive immune cells present in the tumor microenvironment, called Immunoscore, represent an independent prognostic factor for CRC patients, regardless of MSI phenotype [6– 8]. While it is well established that tumor-intrinsic features control the immune response to cancer, we and others have demonstrated the contribution that host tissue-specific factors make to modulating tumor growth and immunity, as well as to response to immunotherapy [9,10]. For instance, using orthotopic mouse models, we have previously shown that kidney and CRC tumors respond poorly to immunotherapy compared to subcutaneous tumors due to organ-specific differences in tumor immune microenvironments [9]. Therefore, a better characterization of the tumor-related immune response in an organ-specific manner could be instrumental for guiding the development of future therapeutics.

Orthotopic xenograft models of CRC established in highly immunocompromised mice recapitulate many features of human pathology and have helped to elucidate several molecular mechanisms involved in CRC progression [11–13]. Nonetheless, human CRC cells xenografted in immunodeficient mice are not exposed to an immune response, which limits their relevance in terms of clinical translation [14]. Therefore, *in vivo* syngeneic orthotopic models are needed to understand the impact of the local immune response during CRC development. Orthotopic implantation refers to the grafting of cells in their original location, thus favoring the generation of an appropriate tumor microenvironment. Transplantation models also allow synchronous growth of implanted tumors in all mice, among other advantages.

In order to study the impact of colon location and the involvement of the immune microenvironment in tumor development, we relied on a pre-clinical immunocompetent orthotopic CRC mouse model. In this model, we used the C57BL/6 (B6)-background MC38 murine CRC cell line [15], recently characterized as a model for hypermutated/MSI CRC [16], that has been genetically engineered to express firefly Luciferase (MC38-fLuc). MC38-fLuc cells were implanted into the caecum (IC) of immunocompetent B6 mice, which allowed us to follow tumor development over time using a bioluminescence camera as well as the dynamic of tumor-infiltrating immune cells by flow cytometry and transcriptomic analyses. Within 2 days, the MC38-fLuc cells developed a growing tumor mass in the colon of all mice, thereby confirming their tumor-forming capacity. Nonetheless, from day 10 onward, we observed two patterns of CRC development in mice with either large lethal colonic tumors and associated metastases (progressive CRC), or spontaneous rejection of tumors (rejecting CRC). This dichotomy in cancer development is colon-specific and associated with a protumor polarization of the immune response in progressive CRC mice and with an antitumor immune microenvironment in rejecting CRC mice. In addition, transcriptomic analysis of CRC tumors at day 3 post-implantation revealed that the two developmental profiles of CRC might be dictated by early immune events.

## Materials and Methods

### Mice and cell lines

Female B6 and BALB/c mice were purchased from Janvier Laboratory (Le Genest-Saint-Isle, France). All experimental protocols were approved by the regional Ethic Committee of Toulouse Biological research Federation (C2EA −01, FRBT) and by the French minister for Higher Education and Research. For the guidelines on animal welfare, we followed the European directive 2010/63/EU.

MC38 parental and firefly-luciferase^+^ (fLuc) cells (B6 background, RRID:CVCL_0A67) and CT26 (BALB/c background, ATCC Cat# CRL-2638, RRID:CVCL_7256) cells were kindly provided by Dr Myriam Capone and Sonia Netzer (ImmunoConcept, CNRS UMR5164, University of Bordeaux, France) and generated as previously described [11]. Cells were cultured at 37 °C and 5 % CO2 in Dulbecco’s Modified Eagle’s Medium (DMEM) (Sigma, Cat#6429) supplemented with 10% fetal bovine serum (FBS) (Gibco, Cat#10270106). Cells were cultured for 2 to 6 passages and tested negative for mycoplasma.

### Tumor implantation

Mice were implanted intra-colon (IC), into the caecum, as previously described [11] [12]. 1×10^6^ or 3×10^6^ viable tumor cells were injected IC sub-serosa. To ensure optimal reproducibility, injections were always performed in the same site of the caecum (Additional file 1: Figure S1A). Subcutaneous (SC) solid tumors were generated by injecting 3×10^6^ viable tumor cells in the right flank of the mouse. Tumor progression was measured using a caliper and mice were euthanized when tumor size reached the ethically defined limit of 250 mm^2^. Intra-hepatic (IH) solid tumors were established by injecting 2.5×10^5^ viable tumor cells in anaesthetized-mice as previously described [9].

### Bioluminescence imaging

*In vivo* MC38-fLuc tumor growth and invasion was monitored, twice a week, using the cooled charge-coupled device camera IVIS Spectrum *in vivo* Imaging System (PerkinElmer), following intraperitoneal (IP) injection of 150 mg/kg of D-Luciferin (Oz Bioscience, Cat#LN10000). Quantitative analyses were performed using IVIS Living Image 4.5.2 software (PerkinElmer; RRID:SCR_014247). Bioluminescent signal intensity was presented as average radiance (photons/sec/cm^2^/sr). For *ex vivo* imaging, mice were administered D-Luciferin, sacrificed and organs of interest were rapidly excised and immersed in 12-well plate with 4.5 mg/mL D-luciferin.

### Tumor processing and flow cytometry

IC tumors were digested for 30 min in DMEM with 1mg/ml Collagenase Type IV, 50 U/ml DNase I Type IV and 100 µg/ml Hyaluronidase type 4 (all from Sigma, Cat#C5138, D5025, H6254) at 37 °C with agitation followed by filtration through a 70 µM cell strainer. For spleen single-cell suspension preparation, spleens were dissociated by filtration through a 70 µM cell strainer. An Ammonium-Chloride-Potassium (ACK) buffer erythrocyte lysis step was then performed. Cells were resuspended in PBS with 2% FBS containing anti-mouse CD16/CD32 antibody (table 1) and 1:1000 Fixable Viability Dye eFluor™ 506 (eBioscience, Cat#65-0866) or Zombie UV™ Fixable Viability Kit (BioLegend, Cat#423108). Staining with primary fluorophore-conjugated antibodies directed against cell surface markers (Additional file 4: Supplementary methods) was performed. Flow cytometric analyses were performed on a LSR II flow cytometer (BD Biosciences) and analyzed using FlowJo software (Treestar, RRID:SCR_008520).

### Statistical analysis

Results are expressed as the mean or median ± standard error of the mean (SEM). All statistical analysis was performed with Prism software (GraphPad Prism, RRID:SCR_002798). The variation in survivals between different groups was analyzed using Log-rank (Mantel-Cox) test. Experiments were analyzed using Mann-Whitney test and P<0.05 is considered significant.

The list of *antibodies used in flow cytometry* and the methods related to *antibody administration for in vivo depletion*, *RNA extraction*, *Luciferase qPCR* and *Nanostring and computational analysis* are provided in Additional file 4: Supplementary methods

## Results

### Colon orthotopic implantation leads to two cancer development profiles

To study the impact of the immune microenvironment on the development of colorectal tumors, we used an orthotopic CRC syngeneic mouse model. Using a microsurgery approach (Additional file 1: Figure S1A), syngeneic colon tumor cells were implanted into the colon (IC) of BALB/c and B6 mice. Following IC injection of CT26 tumor cells, all BALB/c mice developed lethal CRC (Additional file 1: Figure S1B), in accordance with our previous studies [9]. In contrast, following IC implantation of MC38-fLuc tumor cells, two out of five B6 mice survived (Additional file 1: Figure S1B). Since all B6 mice had a CRC tumor at day 6 (Additional file 1: Figure S1C), we hypothesized that the higher survival rate of B6 mice was due to spontaneous rejection of the tumor. To test this hypothesis, we monitored *in vivo* growth of CRC tumors in B6 mice (Additional file 1: Table, n=110) by measuring the bioluminescence emitted by MC38-fLuc cells over time. We previously demonstrated in several orthotopic tumor models that tumor size positively correlates with the bioluminescence of luciferase-transduced tumor cells [9,11]. On days 3 to 6 following IC injection of MC38-fLuc cells, we again found that 100% of B6 mice exhibited solid tumors of comparable size (Fig. 1), thereby confirming efficient orthotopic tumor implantation and initial tumor growth in all mice. However, starting from day 10 after tumor implantation, ~29% of mice developed progressive invasive lethal CRC (progressive group), while ~71% of mice spontaneously rejected the CRC tumors and survived more than 100 days (rejecting group) (Fig. 1, Additional file 1: Table). Accordingly, macroscopic tumors observed in mice with progressive CRC were no longer visible on the caecum of rejecting CRC mice (Fig. 1C). We also noticed the presence of mesenteric lymph node tumor dissemination (Additional file 1: Figure S1D) in mice with progressive CRC. Altogether, our results showed that despite the initial growth of MC38-fLuc tumors in the caecum of all B6 mice, nearly three quarters of them spontaneously rejected the tumor and one quarter of the animals developed progressive, invasive and lethal CRC.

**Figure 1:**
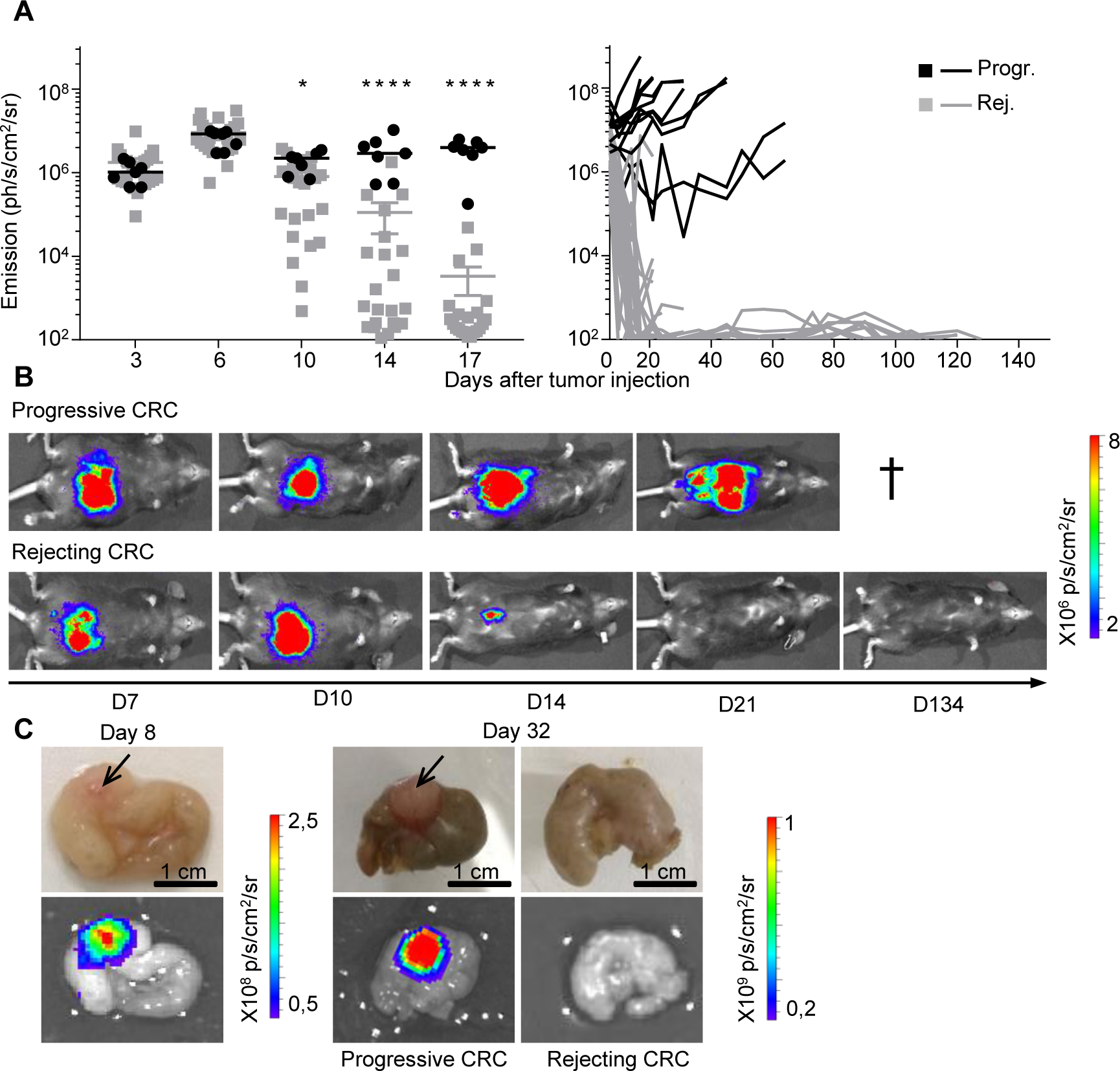
Two profiles of CRC development in immunocompetent B6 mice. **A, B**: Bioluminescence emission monitoring (**A**) and image of one representative mouse per group (**B**) from progressive (Progr.) and rejecting (Rej.) CRC groups following IC-injection with 1×10^6^ MC38-fLuc cells. (Average ±SEM, *n*=36 mice, representative experiment of 3). **C**: *Ex vivo* photograph (upper panel) and corresponding bioluminescent image (bottom panel) of representative caecums from day D8 and D32 MC38-fLuc IC-implanted mice. Arrows indicate tumors on the caecum. *P<0,05, ****P<0,0001

### In rejecting mice tumor cells do not survive in a dormant state but are rather eliminated

All B6 mice that rejected the CRC tumors exhibited an extinction of the bioluminescent signal within 30 days, a delay after which macroscopic tumors were no longer visible. Nonetheless, during the long-term monitoring of CRC progression over a 100-day period, we sometimes detected a weak caecum-localized bioluminescent signal in ~42% (±12% in three independent experiments) of these mice (Fig. 2A and 2B). As bioluminescence detection generally has a negligible background [17,18], we hypothesized that luciferase-expressing tumor-cell variants may have survived in a dormant state in the caecum of tumor-rejecting mice [19]. Adaptive immune cells, in particular CD4^+^ and CD8^+^ T-lymphocytes, are essential for the establishment and maintenance of tumor dormancy [20]. Depletion of T lymphocytes during tumor dormancy has been shown to promote tumor regrowth [20]. Thus, in order to evaluate whether tumor cells in tumor-rejecting mice were dormant, we depleted CD8^+^ and CD4^+^ T-lymphocytes by repeated intraperitoneal injections of anti-CD8 and anti-CD4 antibodies. Despite an effective depletion of CD8^+^ (13% to 3.4% following depletion, P<0.05) and CD4^+^ (18% to 4.7% following depletion, P<0.005) T cells (Additional file 1: Figure 2A and 2B), we did not observe any tumor relapse in these mice (Fig. 2C), suggesting that tumor elimination had occurred. To confirm the elimination of CRC tumors, we measured the presence of luciferase transcripts in colon fragments using qPCR (Additional file 1: Figure S2C). We could detect luciferase-expressing tumor cells in the colons of progressive tumor-bearing mice but not in those of CRC-rejecting mice (Fig. 2D). In addition, luciferase-genomic qPCR (Additional file 1: Figure S2D) confirmed the absence of residual MC38-fLuc cells that may have switched off luciferase gene expression during dormancy (Fig. 2E). Together, these data demonstrate that following initial CRC tumor growth, the group of tumor-rejecting mice spontaneously eliminated MC38-fLuc cancer cells.

**Figure 2:**
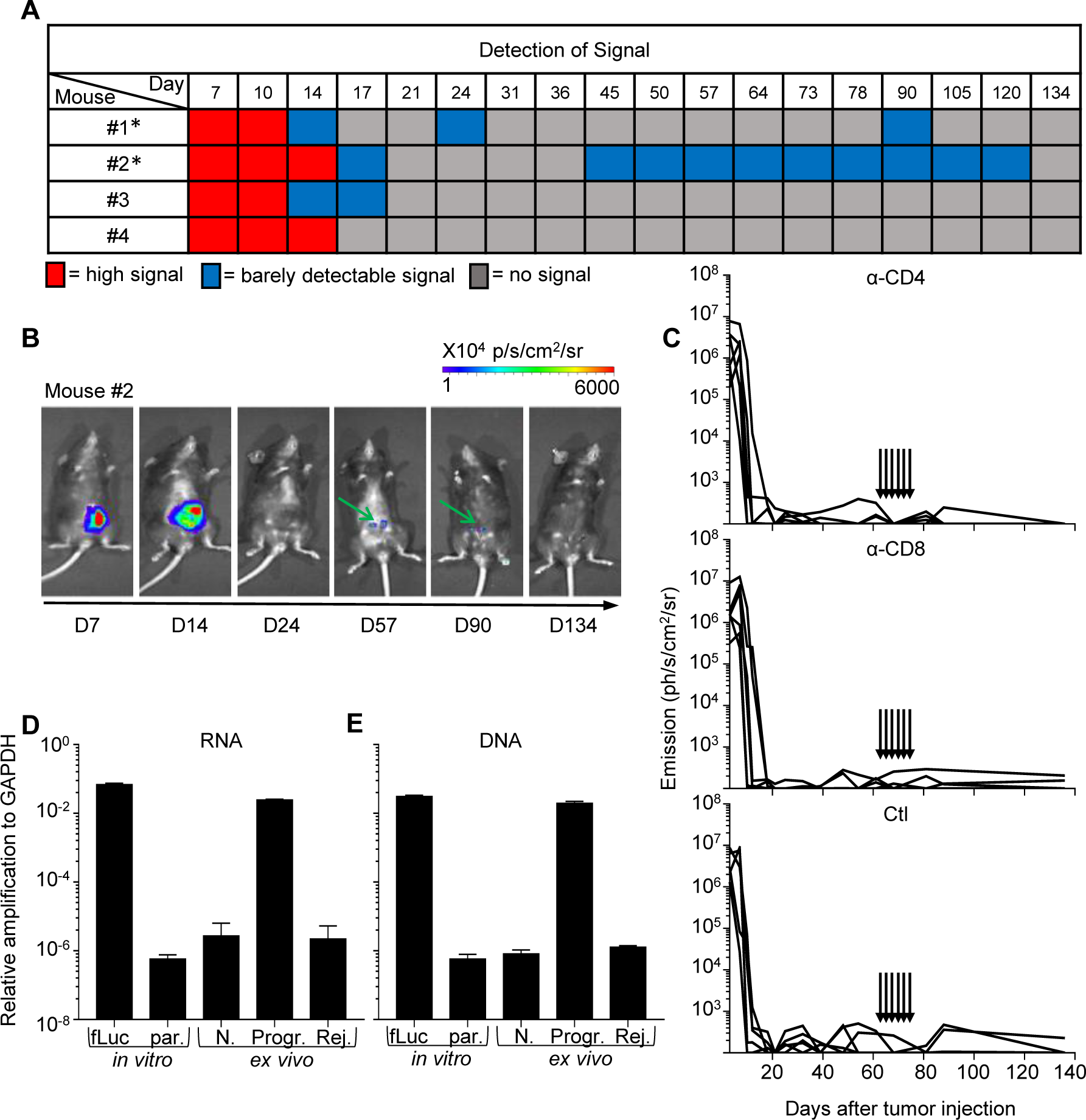
Rejection of MC38-fLuc tumors leads to elimination. A: Bioluminescent signals of 4 representative mice from the rejecting CRC group following IC-implantation of 1×106 MC38-fLuc cells. Mice with * exhibit possible dormancy. B: Bioluminescent images of a representative mouse (#2 from A) possibly exhibiting dormant tumor cells at various time points. Green arrows indicate the weak signal detected at day (D) 57 and 90. C: Bioluminescence emission monitoring in mice from the CRC-rejecting group (following D0 IC injection of 1×106 MC38-fLuc), depleted through injection (indicated by arrows) of anti(α)-CD4 antibody (Ab) (upper panel, n=6), α-CD8 Ab (middle panel, n=6) or control (ctl, bottom panel, n=5). D, E: Relative expression of RNA following RTPCR and qPCR (RNA, D) and amplification of genomic DNA following qPCR (DNA, E) in colons dissected from rejecting-CRC mice (Rej. ex vivo, n=13 with n=6 at D84 and n=7 at D194 post IC MC38-fLuc tumor implantation). Controls (ctl) are represented by in vitro tissue-culture MC38-fLuc (fLuc, positive ctl) and parental (par., negative ctl) cells, ex vivo naïve caecum (N., negative ctl) and D61 IC-tumor bearing mice from progressive-CRC group (Progr., positive ctl) (n=2 to 4). According to negative controls, values below 10-6 are considered background. The relative amplification was normalized to the amplification level of GAPDH.

### CRC fates are specific to the colon immune microenvironment

In order to increase the frequency of progressive CRC-mice, we increased by three folds the number of injected MC38-fLuc cells. Following injection, 50% of the mice still rejected their CRC, demonstrating that the dose of injected cells does not fully explain the two distinct CRC development profiles (Fig. 3A). Besides, both CRC developmental profiles were observed after IC implantation of parental MC38 cells (Additional file 1: Figure S2E), indicating that spontaneous CRC rejection is not related to the potential immunogenicity of luciferase in MC38-fLuc cells. In addition, no tumor rejection was observed after injection of MC38-fLuc cells in another anatomical (non-orthotopic) location in the mice, such as in the liver (IH, Fig. 3B) or subcutaneously (SC, Fig. 3C). We then evaluated the contribution of the immune response to CRC tumors rejection by injecting MC38-fLuc cells into the caecum of fully immunodeficient NOD/SCID mice. We observed that all mice developed lethal CRC, characterized by larger tumors than those observed in immunocompetent B6 mice (average emission 6,9×10^7^ (NOD/SCID) versus (vs) 2,9×10^6^ (B6) ph/s/cm²/sr at day 9, P<0.0001, Fig. 3D and 3E). Accordingly, NOD/SCID mice died more rapidly following tumor implantation (before day 15) than B6 animals, and never rejected CRC tumors (Fig. 3E), underlining the importance of the immune response in CRC outcome. We performed a similar experiment in nude mice, which only lack adaptive immunity. After tumor implantation, nude mice developed lethal CRC and died within 15 days (Fig. 3D and 3E), confirming the key contribution of the adaptive immune response to tumor rejection. Nude mice developed larger tumors than progressive-B6 and NOD/SCID mice (nude average emission 4,7 x10^8^ ph/s/cm²/sr at day 9, Fig. 3D), suggesting the innate immune response may facilitate CRC tumor progression. Taken together, these data demonstrated that the colon-specific immune microenvironment is responsible for tumor rejection in the majority of B6 mice. We next asked whether this immune microenvironment may have led to a clonal selection of “immune-resistant progressive” tumor cells in the CRC-progressive mice *vs*. “immune-sensitive rejecting” tumor cells in the CRC-rejecting mice. To address this question we harvested and cultured MC38-fLuc tumor cells from both groups of mice at day 14 after the initial tumor implantation and re-implanted them IC in recipient naïve B6 mice (Additional file 1: Figure S3A). Re-implantation of MC38-fLuc derived from both CRC-progressive and CRC-rejecting mice led each to the development of two CRC profiles in naïve recipient mice (Additional file 1: Figure S3B), indicating that *in vivo* clonal selection is not the main determinant regulating progression or rejection of tumor cells in our model.

**Figure 3:**
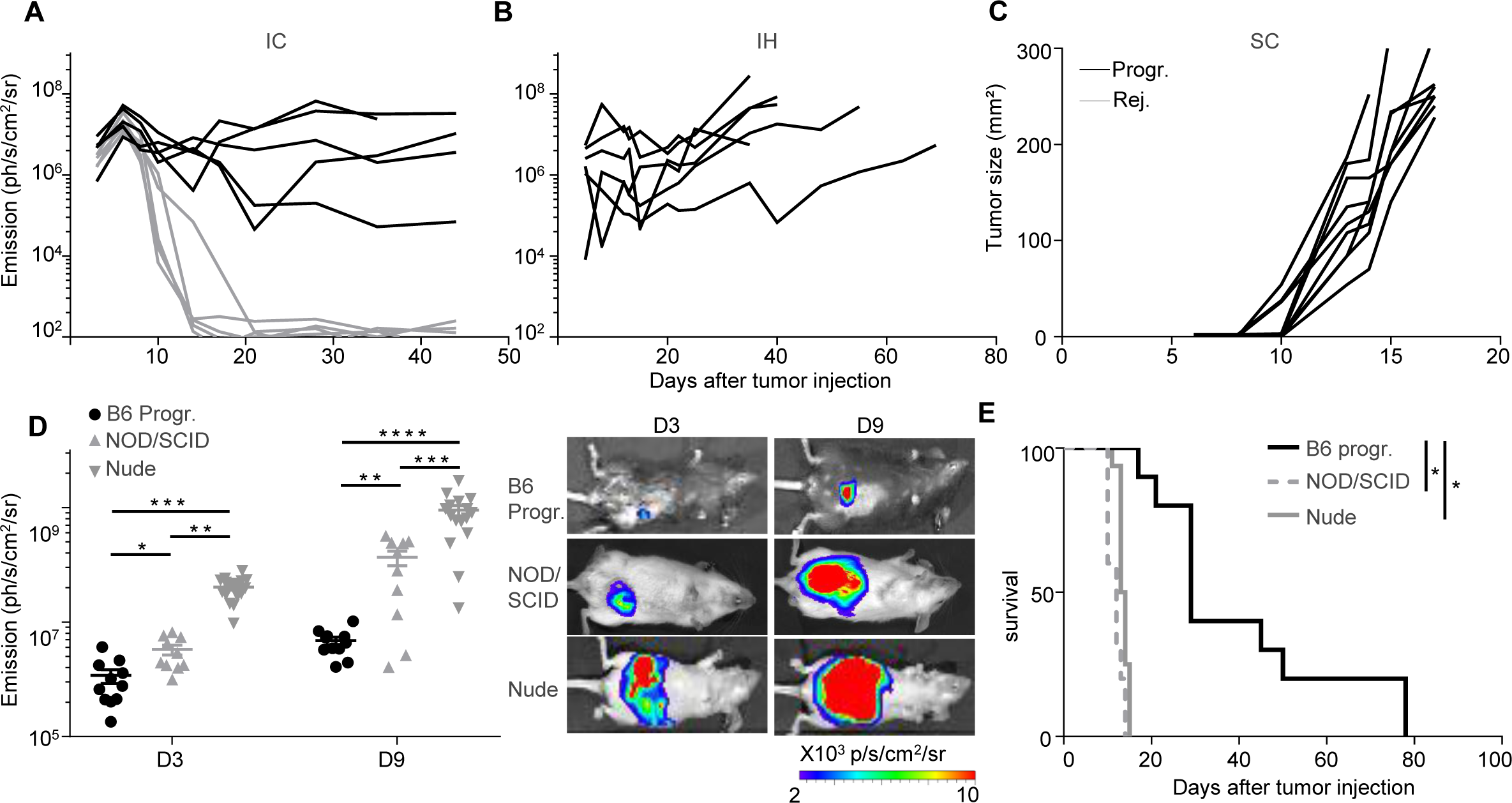
An immune-colon dependent effect generates the two CRC development profiles. **A to C**: Tumor growth monitoring through bioluminescence emission imaging (**A, B**) and caliper measurement (**C**) of 3×10^6^ IC-injected MC38-fLuc cells *(n=10*) (**A**), 2,5×10^5^ IH-injected MC38-fLuc cells (*n*=6) (**B**) and 3 x10^6^ SC-injected MC38-fLuc cells (*n*=9). **D, E**: Bioluminescence emission imaging (Average±SEM) (**D**) and survival (**E**) of 1×10^6^ MC38-fLuc IC-injected in B6 mice (depicted from progressive CRC group, *n*=11), NOD/SCID mice (*n*=10) and nude mice (*n*=16). (**D**) Graph of bioluminescent emission (left panel) and representative photos of one mouse per group (rigth panels) at day (D)3 and D9. *P<0,05, **P<0,005; ***P<0,0005, ****P<0,0001.

### Tumor fates correlate with suppressor/effector immune microenvironments

In order to investigate the immune microenvironment of tumors from the two CRC profile groups, we used flow cytometry to examine the phenotype and frequency of infiltrating leukocytes in primary colon tumors of CRC-rejecting *vs*. CRC-progressive mice at 9, 20 and 29 days post-tumor implantation (Fig. 4A and 4B). As the “group separation” is not fully resolved at day 9 (Fig. 1A), we measured decrease or increase in colon tumor size compared to day 6 (using bioluminescence) to differentiate the mice belonging to CRC-rejecting or CRC-progressive groups.

**Figure 4:**
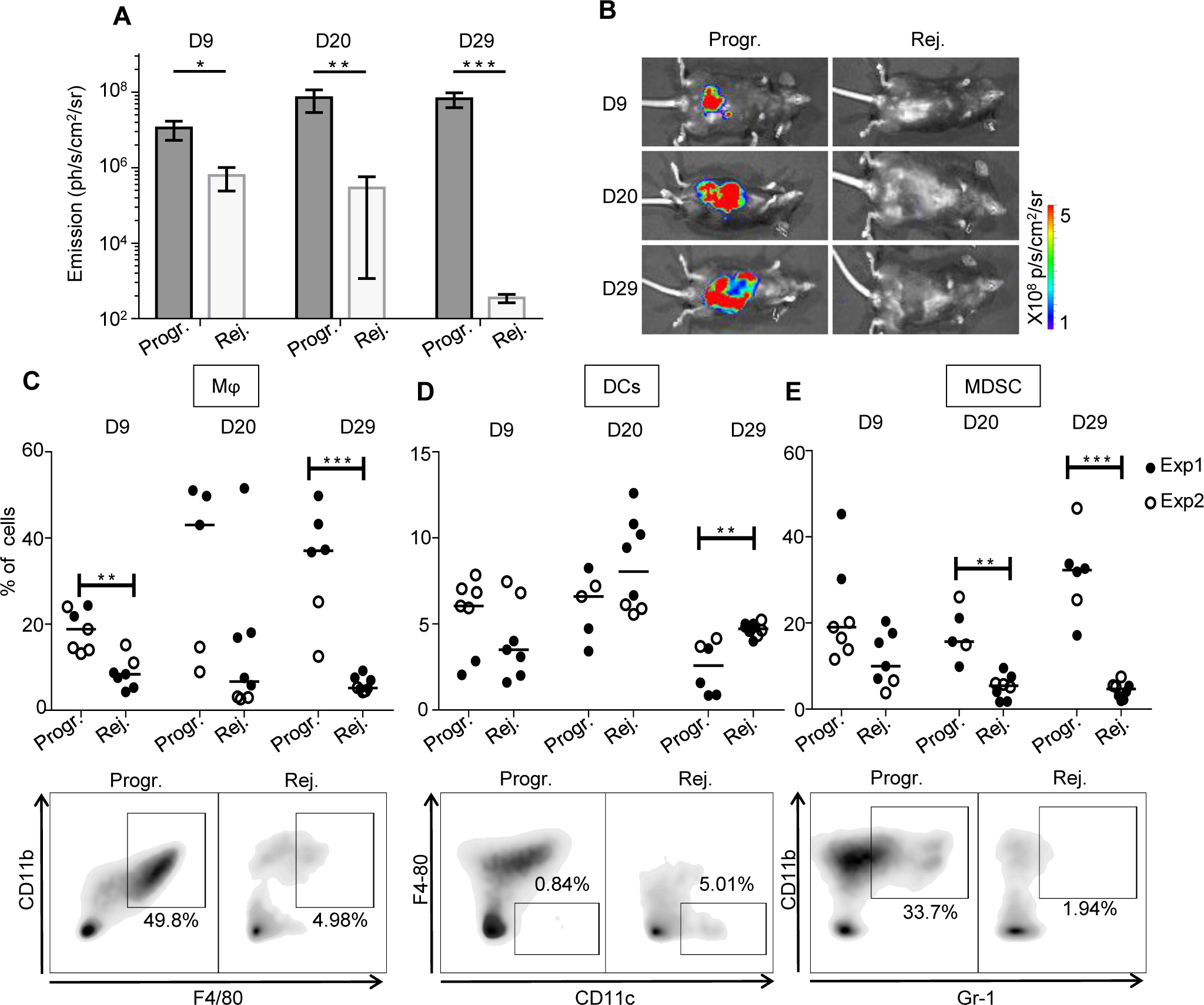
The immunosuppressive myeloid cell-related microenvironment is characterized in progressive CRC mice. **A,B**: Bioluminescent emission measurement (**A**) and representative photo of one mouse per group used for FACS analyses (**B**) at day (D) 9, 20 and 29 following IC-implantation of 1×10^6^ MC38-fLuc cells at D0. (Average ±SEM, *n*=6-7 mice, 2 pooled experiments). **C to E**: Quantitative data (upper panels) and D29 representative FACS dot plot analyses (lower panels) of progressive or rejecting tumors infiltrating macrophages (Mϕ, gated as Gr1^−^, F4/80^+^, CD11b^+^) (**C**), dendritic cells (DCs, gated as Gr1^−^, CD11c^+^, F4/80^−^) (**D**) myeloid-derived suppressor cells (MDSC, gated as Gr-1^+^, CD11b^+^) (**E**) at D9, 20 and 29. Each dot correspond to a single mouse with median from two independent experiments. Percentages are expressed on CD45.2^+^ live cells. *P<0,05, **P<0,01 and ***P<0,001. Progr.= progressive CRC, Rej.= Rejecting CRC.

Within the myeloid cell compartment, we observed increased infiltration of F4/80^+^/CD11b^+^ macrophages in tumors of CRC-progressive mice compared to CRC-rejecting animals (Fig. 4C). Macrophages infiltrating advanced tumors are known to exert mainly immunosuppressive functions, in particular when polarized toward an anti-inflammatory type-2 macrophage (M2) phenotype [21]. While tumor-infiltrating macrophages from progressive CRC mice expressed low levels of the M2-related marker CD206 [21] on day 9 post-tumor implantation, these levels were significantly higher than in rejecting mice CRC tumors. CD206 expression increased substantially by days 20 and 29 although there was no longer a significant difference between the two CRC profiles (Additional file 1: Figure S4A). In addition, high levels of the type-1 (pro-inflammatory) macrophage (M1) related marker CD80 [21] were detected on macrophages from progressive CRC tumors (Additional file 1: Figure S4C), implying that the CD206 and CD80 markers may not be sufficient or appropriate markers to discriminate M2 and M1 polarization status in colon tumors. Tumor infiltrating macrophages from progressive CRC mice strongly expressed MHCII, CX3CR1, and to a lesser extent CD11c, while they were negative for Ly6C and CD103 (Additional file 1: Figure S4C). Regarding other myeloid cells, we observed a higher infiltration of total CD11c^+^ dendritic cells (DC) in the rejecting group on day 29 after tumor injection but not at previous time points (Fig. 4D). No significant difference in CD206 expression in DC between the two groups of mice (Additional file 1: Figure S4B). Finally, we observed that during CRC progression, progressive tumors contained a higher infiltrate of myeloid-derived suppressor cells (MDSC), a major immunosuppressive cell subset in tumors [22], compared to rejecting tumors (Fig. 4E).

CD8^+^ T cells, which are involved in adaptive immune responses, highly infiltrated rejecting tumors. This was particularly noticeable at day 29 (Fig. 5A). CD8^+^ T cells infiltrating progressive tumors expressed significantly higher levels of the immunosuppressive checkpoint PD-1 (Fig. 5D). PD-1 expression was also increased on the surface of CD4^+^ T cells that infiltrated progressive tumors (Fig. 5E), suggesting a stronger exhaustion status of both CD4 and CD8 T-cell populations in progressive compared to rejecting CRCs. Although the total CD4 T-cell infiltrate was comparable in the two tumor profiles (Fig. 5B), we found a significant increase in the percentage of immunosuppressive regulatory CD4^+^ T cells (Treg) during tumor development in progressive compared to rejecting colon tumors (Fig. 5C). We also observed that at day 20, Treg cells from progressive tumors expressed high levels of PD-1 (Fig. 5F). Taken together, these results demonstrate that our orthotropic CRC tumor model exhibits two patterns of CRC-targeted immune response, cancer repressing and cancer promoting. The immune microenvironment of the tumors appears to be biased towards the antitumor axis of the immune response in the rejecting CRC mice, *vs*. the protumor axis in the progressive CRC mice.

**Figure 5:**
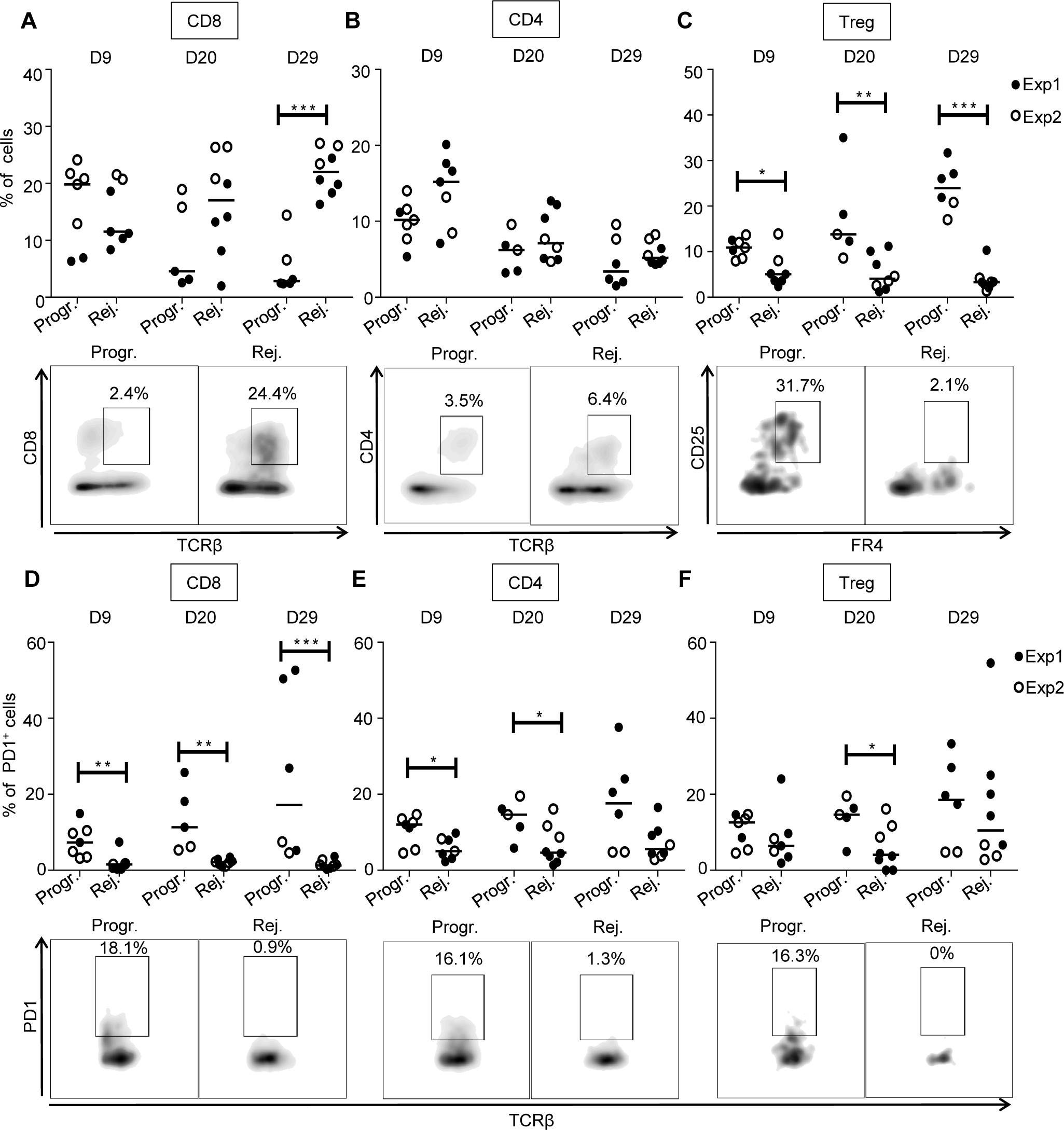
High CD8 T cells and low Treg infiltration in rejecting CRC tumors. Quantitative data (upper panels) and D29 (**A, B, C**) or D20 (**D, E, F**) representative FACS dot plot analyses (lower panels) of tumor infiltrating CD8^+^ T lymphocytes (CD8, gated as TCRβ^+^, CD8^+^) (**A**), CD4^+^ T lymphocytes (CD4, gated as TCRβ^+^, CD4^+^) (**B**), regulatory T lymphocytes (Treg, gated as TCRβ^+^, CD4^+^, CD25^+^, folate receptor (FR)4^+^) (**C**) as well as PD1 expression on CD8^+^ T lymphocytes (**D**), CD4^+^ T lymphocytes (**E**) and regulatory T lymphocytes (**F**) at 9, 20 and 29 days, as depicted, following IC-injection of 1×10^6^ MC38-fLuc cells. Each dot correspond to a single mouse with median from two independent experiments. Percentages are expressed on CD45.2^+^ live cells. *P<0,05, **P<0,01 and ***P<0,001. Progr.= progressive CRC, Rej.= Rejecting CRC.

### Early polarization of the colon immune microenvironment determines tumor fate

We hypothesized that early events may be responsible for the dual immune microenvironments characterizing the two opposite profiles of cancer development in our mouse model. To test this assumption, we implanted MC38-fLuc IC in B6 mice (n=19), harvested solid tumors on day 3 post-implantation (Additional file 1: Figure S5A) and measured the expression of 770 cancer-immune related-genes, in the whole tumors, using a quantitative and multiplex method referred to as Nanostring nCounter Gene Expression Assay. Based on either the most expressed genes (P>0.005, 67 genes, Fig. 6) or on the significantly expressed genes (P>0.05, 187 genes, Additional file 1: Figure S5B and Additional file 3), we observed that mouse tumor samples clustered into two distinct groups (52% dark-red group and 48% blue group represented on the heat maps). The first cluster of mouse tumors (blue group) exhibit a high expression of genes (10 out of 19 highest up-regulated genes) previously associated with Treg cells including *CD4*, *CD247*, *Irgm2*, *Herc6*, *Ccl17*, *Tgfbr2*, *St6gal1*, *Nt5e, Rora* (Fig. 6A and 6B) as well as *Foxp3*, *Maf, Ccl19, Ccl21* (Additional file 1 and 3: Figure S5B) [23–26]. We also found high expression of CD103 (*Itgae*), a marker of gut resident T cells including Treg (Fig. 6A and 6B) [25]. This signature, together with the high expression of *Tgfb2* and *Lag3* suggested that tumors from the blue group are skewed toward a tolerogenic, likely protumor microenvironment (Additional file 1: Figure S5B and additional file 3). In contrast, tumors belonging to the dark-red group displayed a robust pro-inflammatory signature as demonstrated by the strong expression of genes related to the inflammasome (*Nlrp3*, *Il1b, IL1a* in Fig. 6A and 6C), to inflammatory cytokines signaling (*Il1rl1*, *Il1r2* in Fig. 6A and 6C; *Tnf*, *Il6*, *Il23r* in Additional file 1: Figure S5B and additional file 3) as well as to inflammation signaling (*Cebpb*, *Mefv* in Fig. 6A and 6C; *Lyn*, *ptgs2*, *Sbno2* in Additional file 1: Figure S5B and additional file 3) [27,28]. Accordingly, we observed a high gene expression of markers related to pro-inflammatory innate immune cells including *Trem1*, *Clec7a*, *Csf2rb, Nos2*, *Fcgr3*, *CD86* (Fig. 6A and 6C) and *Csf3r*, *CD87, Fcgr2b*, *CD47* (Additional file 1: Figure S5B and additional file 3) as well as a high expression of genes related to myeloid cells recruitment such as *Ccl2*, *Ccl3*, *Ccl7*, *Ccr1*, *Cxcl1*, *Cxcl2*, *Selplg*, *Cxcl5* (Fig. 6A and 6C). Most of the myeloid compartment up-regulated genes were related to monocyte/macrophage populations with high expression of *Cd80*, *VegfA* (Fig. 6A and 6C) and *CD14*, *CD68* (Additional file 1: Figure S5B and additional file 3) but we also detected an increased expression of neutrophil markers (*Ppbp*, *Ncf4*, *Cxcr2*, Fig.6A and 6C) [29–32]. Finally, in the dark-red group of tumors, we observed the upregulation of *Ifnar1* gene reflecting the initiation of a cytotoxic immune response [33] related to T lymphocytes and NK cells recruitment and activation (*lcp1*, *Sell*, *Tnfsf14* in Fig. 6A and 6C; *Prf1*, *Itgb2, CD244*, *Klra17*, *Klra2*, *Il21*, *CD53* in Additional file 1: Figure S5B and additional file 3) and suggesting an antitumor polarization of the immune microenvironment [32,34]. To summarize, at day 3 post tumor-implantation, half the mice (blue group) present an IC-tumor immune microenvironment biased toward an immunosuppressive, protumor profile while the other half (dark-red group) have an IC-tumor microenvironment indicating the initiation of a proinflammatory, hence antitumor, response.

**Figure 6:**
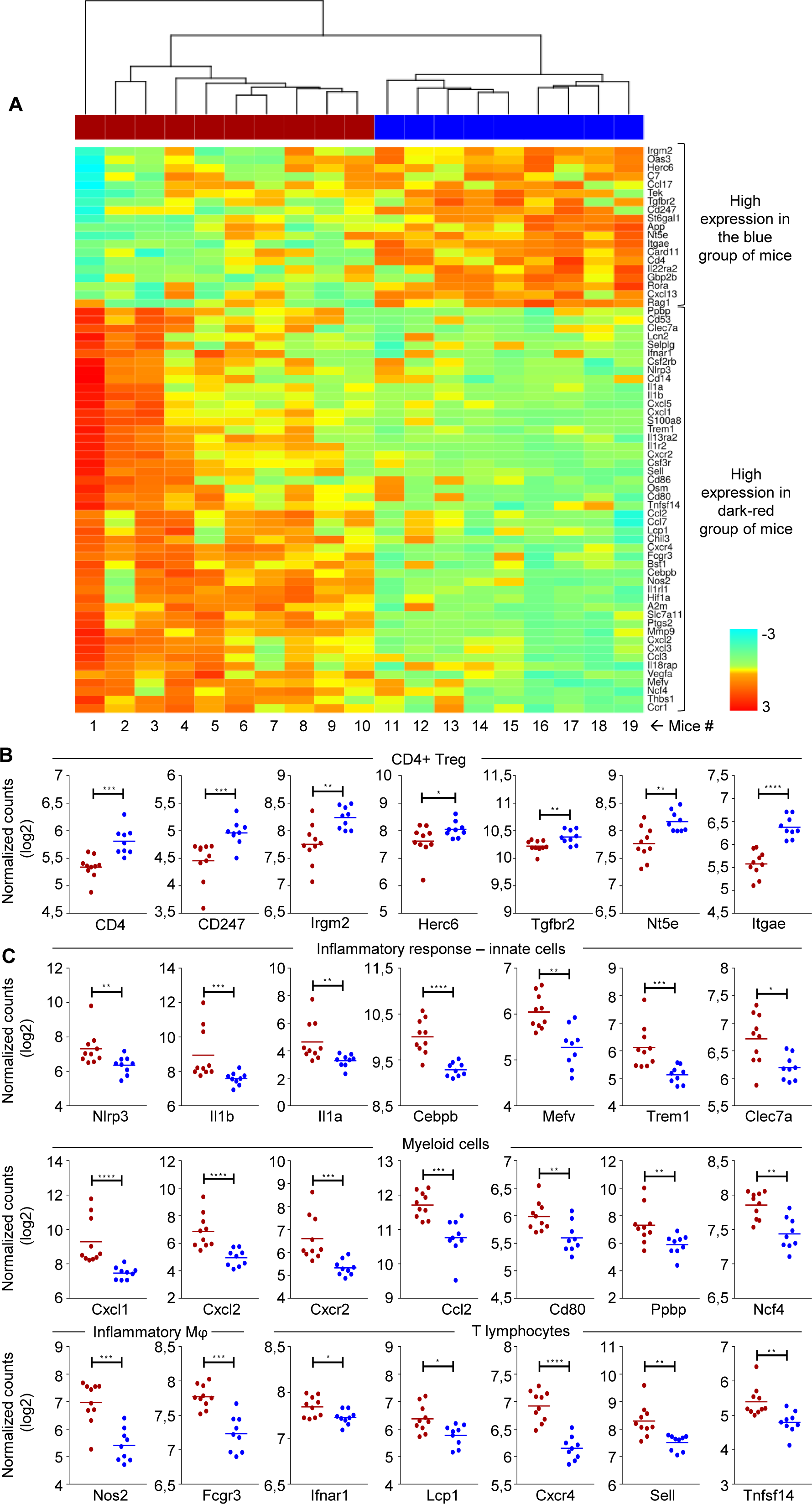
Early detection of opposing immune microenvironments in CRC tumors. (**A**) Heat map representing the Z-score normalized expression of genes, measured by Nanostring, that are up- or down-regulated in whole CRC tumors extracted at 3 days following IC implantation of 1×10^6^ MC38-fLuc cells in B6 mice. Only the genes whose expression varies significantly (p-value< 0.005 based on ANOVA test) are represented on the heat map. Based on the differences of gene expression, two groups of tumors (samples 1 to 10 in dark-red and samples 11 to 19 in blue) can be distinguished on the heat map. (**B, C**) Dot plots representing the average and individual tumor expression of selected genes (normalized counts in log 2) analyzed by Nanostring technology. Dark-red and blue dots correspond to tumors belonging to the dark-red or blue groups respectively. Selected genes highly expressed in (**B**) blue group (CD4+ Treg pattern) and (**C**) dark-red group of tumors (Inflammatory response-innate cells, myeloid cells, inflammatory macrophages (Mφ) and T lymphocytes are depicted. *P<0,05, **P<0.005, ***P<0.0005, ****P<0,0001

## Discussion

Most preclinical mouse models investigating the immune composition of CRC tumors are based on the implantation of colon tumor cell lines under the skin [35,36]. Nevertheless, our past work has revealed that the skin microenvironment does not accurately reproduce the microenvironment of organs from which tumor cells originate [9]. The colon tumor model described here relies on transplanting tumor cells in their tissue of origin, *i.e.* the colon, in order to reconstitute an appropriate immune microenvironment. Using this methodology, we previously demonstrated that the orthotopic injection of the syngeneic CT26 colon tumor cells into BALB/c mice led to the systematic development of lethal CRC disease associated with an immune microenvironment that was systematically protumor biased [9]. Here, we found that orthotopic injection of MC38 colon tumor cells into the caecum of B6 mice gave rise to two spontaneous and opposing immune microenvironments in the colon, ultimately leading to either the elimination of tumors or the promotion of cancer development.

Using MC38-fLuc cells and an *in vivo* bioluminescence monitoring approach, we demonstrated that tumor implantation and growth were identical in all mice at the early stage of disease development (before day 10). Nonetheless, from day 10, tumor persisted and progressed to lethal CRC in only 25-35% of mice, whereas the remaining 65-75% of animals spontaneously rejected CRC tumors. Other research groups have previously performed similar orthotopic-implantation of parental MC38 cells or MC38-fLuc in the colon of B6 mice [37– 39]. These studies have reported a rather low tumor incidence (25% to 40%) between 4 to 6 weeks post-implantation, which was explained by a poor tumor intake. However, in the absence of longitudinal monitoring of tumor growth by bioluminescence detection, these authors were unable to observe that tumor cell implantation was similar in all mice and most probably missed the rejection phase that we detected, in 70% of mice, during the second week post-implantation (*see Fig. 1*). In addition, we initially optimized CRC development by injecting 1×10^6^ MC38 cells, which represents a lower tumor-cell dose than used in the above cited studies (i.e. 2×10^6^ MC38 cells) [37–39]. Injecting three times more cells (3×10^6^), to optimize the chances of tumor implantation, did not dramatically increase the percentage of mice developing a progressive CRC profile, thus implying that the number of injected cancer cells has a negligible impact on the two CRC developmental profiles. Although rarely observed in B6 mice [40], increased immunogenicity linked to the expression of luciferase in tumor cells was previously described in other tumor mouse models [41]. However, since the outcome of orthotopic parental MC38 tumors was the same as that of MC38-fLuc IC-tumors, we concluded that luciferase immunogenicity could not explain the CRC rejection profile.

The immune response plays a decisive role in determining the outcome of CRC in our mouse model. Indeed, we never observed tumor rejection in the absence of functional innate and/or adaptive immune components. In line with our previous study highlighting the decisive role of anatomical location in shaping tumor immunity [9], the rejection of MC38 tumors has not been observed when MC38 cells were implanted in other locations than the colon (i.e. under the skin or in the liver) [42]. Altogether, our data underline a critical role for colon-specific determinants in regulating the polarization of the tumor immune response, leading either to the rejection or the progression of CRC. Among these determinants, the microbiota was recently shown to be a key component involved in the polarization of the colon-specific immune response during CRC and its composition may vary between mice from various providers [43]. Nonetheless, differences in microbiota may not explain the different CRC development profiles in our mouse model as all animals used in our experiments come from the same provider and were littermates. Although one could propose that variation in the overall microbiota composition may appear after the arrival of mice in our animal facility, we never observed any “box-effect” that would lend support to such a hypothesis. Besides, given the key contribution of immunity to CRC tumor rejection in B6 mice, we hypothesized that an *in vivo* clonal selection may occur in the colon of mice leading to the enrichment of “immune-resistant” tumor cell clones, able to escape immunity, in the CRC-progressive group *vs.* “immune-sensitive” tumor cell clones in the CRC-rejecting group [16]. However, we found that regardless of their origin, tumor cells behaved similarly and generated both the progressive and rejecting CRC development profiles after re-implantation in naïve B6 mice. These data suggest that clonal selection may not have occurred or does not fully explain the capacity of tumors to grow in CRC-progressive mice.

We distinguished two opposing immune microenvironments with either a predominant protumor polarization in a progressive CRC tumor microenvironment or an antitumor polarization in a rejecting CRC tumor microenvironment. In line with other studies on colonic macrophages [44–46], we identified F4/80^+^/ CD11b^+^/ MHCII^+^/ CX3CR1^+^/CD103^−^/ Ly6c^−^ colon tumor-associated macrophages (TAM) as the most abundant immune subset. They have previously been characterized as high IL-10 producers [44], indicating their immunosuppressive potential, and they mainly infiltrated progressive tumors. A previous study, using a MC38 orthotopic model, has demonstrated the critical role of TAM in CRC development through the remodeling of the extracellular matrix (ECM) composition and structure [46]. The high infiltration of CD11c^−^/Gr1^+^/CD11b^+^ cells observed in tumor-progressive mice, which likely correspond to the typical immunosuppressive MDSC found in the tumor microenvironment [22], supports an immunosuppressive tumor network. As shown by others, we believe that a significant proportion of monocyte-derived MDSC, skewed by the surrounding microenvironment, will rapidly differentiate into potentially immunosuppressive TAM [22]. The boosted CRC tumor growth observed in nude mice highlighted the importance of innate cells (including TAM and MDSC) in sustaining tumor development. Indeed, in addition to suppressing the antitumor immune response, myeloid cells can produce angiogenic factors and cytokines that can remodel ECM, facilitate tumor angiogenesis and growth [46] [47]. Finally, the high tumor-infiltration by Treg in CRC tumor-progressive mice is consistent with the immunosuppressive immune signature of the microenvironment. Additional phenotypic and functional experiments will help further characterization of the complexity of the immune regulatory components and confirm the immunosuppressive functions of MDSC and Treg in the tumors of progressive mice. In contrast, tumors spontaneously rejected from B6 mice exhibited very little infiltration with immunosuppressive Treg, TAM and MDSC but were highly infiltrated with CD8^+^ T cells, key antitumor effectors in CRC [6]. These CD8^+^ T cells express low levels of the immune checkpoint PD-1 suggesting that their antitumor functions are less inhibited compared to progressive tumors-infiltrating CD8^+^ T cells [48].

While two opposing immune response profiles, which we characterized from day 9 tumors, possibly explain the eventual rejection *vs.* progression of CRC tumors in B6 mice, we also questioned the origin of these divergent immune microenvironments. We performed transcriptomic analysis of CRC tumors at day 3, representing an early time-point after tumor implantation. We found that two opposing immune microenvironments can already be distinguished with half the mice exhibiting a dominant Treg signature and the other half presenting an inflammatory innate immune response signature. In the colon, Treg represent a high proportion of CD4^+^ T lymphocytes (up to 30%) and play a central role in regulating immune responses against commensal microorganisms and dietary antigens [49,50]. Thus, the Treg signature observed in half the mice, at day 3 post-tumor implantation, possibly reflects a preexisting immunosuppressive microenvironment that favors tumor growth. In the other half of mice, a dominant inflammatory immune response was initiated likely by tumor cells. The local disruption of the initially tolerogenic colon microenvironment is outlined by the production of chemoattractant factors and the recruitment of inflammatory innate components and cell populations (inflammatory monocytes, macrophages, NK cells), required for the initiation of an effector antitumor immune response and ultimately the rejection of IC tumors. It remains uncertain why such inflammation potentially leads to rejection in the colon but not in other organs (i.e. skin and liver) in which similar tumor implantations were carried out. The 30/70% proportions described from day 9 do not yet seem to be fully established at day 3. We postulate that mice 11 to 15 (Fig. 1A, blue group) may represent intermediate individuals with intermediate expression of some Treg-related genes (e.g. *Irgm2*) and high expression of some inflammation-related genes (e.g. *Mefv*, *Ncf4*). Eventually, most likely before day 9, the inflammatory/cytotoxic antitumor microenvironment may become dominant in 78% of mice (mice number 1 to 14) while 22% of the remaining mice may maintain a protumor, Treg-associated microenvironment, as confirmed by our flow cytometry analyses from day 9 (*see Fig. 5C*).

## Conclusions

Our study demonstrates the spontaneous and early occurrence of opposing immune polarization phenotypes in CRC tumors in a pre-clinical mouse model. During CRC progression, we evidenced a protumor immunosuppressive immune microenvironment in 25% of the animals and an antitumor immune microenvironment in 65% of the animals. Based on recent work [6], an international consortium of 14 centers in 13 countries validated that high immunoscore patients (who showed a high density of cytotoxic tumor-infiltrating CD8^+^ T cells) had the lowest risk of CRC recurrence at 5 years [7]. These analyzes revealed that 22% of the patients had a low immunoscore while 78% had an intermediate-to-high immunoscore [7], similar to the 30% protumor vs 70% antitumor signature and CRC development profile proportions seen in our MC38 CRC orthotopic C57BL/6 mouse model. Our model reveals the intrinsic potential of the colon microenvironment to become polarized towards the non-immunosuppressive antitumor axis of the immune response, characterized by highly CD8^+^ T-cell infiltrated tumors, and may therefore facilitate the study of the mechanisms underlying the CRC immune response, as well assessment of potential immunotherapeutic interventions.

## Additional file 1

**Figure S1**. Development of orthotopic CRC model in B6 mice. A: Schematic representation of the IC injection procedure. Red dot represents the injection spot. B: Survival of BALB/c mice (*n*= 5) and B6 mice (*n*= 5) following IC injection with respectively 1×10^6^ CT26 cells or 1×10^6^ MC38-fLuc cells. C: Day (D)6 bioluminescent emission of the IC injected B6 mice. D: *Ex vivo* photograph (upper panel) and corresponding bioluminescent image (bottom panel) of representative mesenteric lymph nodes from D16 IC-tumor bearing mice from progressive or rejecting CRC groups. Arrows indicate mesenteric lymph node. *, P<0,05. **Figure S2.** Tumor dormancy evaluation. A, B: Flow cytometry analysis (left panels) and representative plots (right panels) of CD4^+^ (A) and CD8^+^ (B) T cells of viable leukocytes from naïve B6 mice spleen following *in vivo* depletion by 6 IP injections of α-CD4 (A) and α-CD8 (B) Ab respectively and PBS as control (ctl) (A and B). Mice were sacrificed at day 12 and splenocytes were stained with Abs against CD45.2, TCRβ, CD4 and CD8. (Average ± SEM, *n*=3). C, D: Range of detection of RNA relative expression following RTPCR and qPCR (RNA, C) and amplification of genomic DNA following qPCR (DNA, D) for a lysed mix of naive B6 colon with a range (from 4 to 1×10^5^ cells) of *in vitro* tissue-cultured MC38-fLuc cells. Negative controls are represented by *in vitro* tissue-cultured parental MC38 (par.) and naïve B6 colon (N.). Positive controls are represented by *in vitro* tissue-cultured MC38-fLuc (fLuc). (E) Proportion of rejecting CRC (Rej.) and progressive CRC (Progr.) mice injected IC with 1×10^6^ of MC38 parental cell line. At day 25, mice were dissected and examined by eye for the presence (Progr.) or absence (Rej.) of CRC tumors. **Figure S3.** Clonal selection is not sufficient to explain the progressive and rejecting CRC profiles. A: Outline of the cross-implantation experiment of ex vivo *immune-resistant progressive* (Progr.) or *immune-sensitive rejecting* (Rej.) MC38-fLuc cells isolated from day (D)14 IC tumor-bearing, respectively progressive or rejecting, CRC donor mice (initially IC implanted with 1×10^6^ MC38-fLuc cells). Following 3 weeks (wks) of *in vitro* tissue culture, 1×10^6^ of *progressive* and *rejecting* cells were separately re-injected in recipients naive B6 mice (*n*=18 mice per group) and monitored for CRC development. B: Proportion of progressive and rejecting CRC recipient mice following implantation of *progressive* cells (top panel) or *rejecting* cells (bottom panel). **Figure S4.** Surface markers expression on tumor-infiltrating macrophages and DCs. A, B: FACS analysis for the expression of CD206 on tumor-infiltrating macrophages (Mϕ, gated as Gr1^−^, F4-80^+^, CD11b^+^) (A) and dendritic cells (DCs, gated as Gr1^−^, CD11c^+^, F4-80^−^) (B) at different 9, 20 and 29 days following IC injection of 1 × 10^6^ MC38-fLuc^+^ cells (*n*=6 to 8 mice). Each dot corresponds to a single mouse with median from two independent experiments (Exp1 and Exp2). **, P< 0.01. Progr.= progressive CRC, Rej.= Rejecting CRC. C: FACS analysis for depicted markers expressed at day 17 by progressive CRC tumor-infiltrating macrophages (Mϕ, gated as CD64^+^, F4/80^+^, CD11b^+^). **Figure S5:** Nanostring analysis of CRC tumors harvested on day 3 after IC implantation of 1.10^6^ MC38-fLuc cells in B6 mice (n=19). (A) Bioluminescent representative images at 2 days post-tumor implantation. (B) Heat map representing Z-score normalized expression of genes up-or down-regulated in D3 tumors following Nanostring analysis of whole tumor tissue. Only the genes whose expression varies significantly (p-value< 0.05 based on ANOVA test) are represented on the heat map. Based on the differences of gene expression, two groups of tumors (samples 1 to 10 in dark-red and samples 11 to 19 in blue) can be distinguished on the heat map. Selected genes are depicted, on the right side, for corresponding line on the heat map.

## Additional file 2

**Table: Proportion of MC38 IC-implanted mice** Numbers and percentages of mice exhibiting a progression or rejection of their MC38-fLuc tumors in three independent experiments (total *n*=110).

## Additional file 3

**Gene expression changes in B6 IC-tumors.** Transcripts differentially expressed with P<0.05 (ANOVA test) are listed for the 19 IC-tumors of B6 mice implanted with 1.10^6^ MC38-fLuc cells at D0 and harvested at D3. Differential expression was determined within the Nanostring PanCancer Immune profiling panel.

## Additional file 4

**Supplemental Methods**

## List of abbreviations

α: anti
ACK: ammonium-chloride-potassium
B6: C57BL/6
CRC: colorectal cancer
DC: dendritic cell
DMEM: dulbecco’s modified eagle’s medium
ECM: extracellular matrix
FBS: fetal bovine serum
IC: intra-colon
fLuc: firefly luciferase
IH: intra-hepatic
IP: intra-peritoneal
M1: type 1 macrophages
M2: type 2 macrophages
MDSC: myeloid-derived suppressor cells
MMR: DNA mismatch repair system
MSI: microsatellite instability high
NK: natural killer
q: quantitative
Treg: regulatory CD4^+^ T cells
SC: subcutaneous
SEM: standard error of the mean
TAM: tumor associated macrophages

## Declarations

### Ethic approval

All experimental protocols were approved by the regional Ethic Committee of Toulouse Biological research Federation (C2EA - 01, FRBT) and by the French minister for Higher Education and Research.

### Consent for publication

Not applicable.

### Availability of data and material

Data used and analyzed during this study are included in this published article (and its additional files).

## Authors’contributions

GT, AFT, VF, FLV, CV, CD performed experiments; GT, AFT, FLV, MT, MA, YR and CD designed the experiments and analyzed the results; AFT, MA, ON, NV, YR, CD provided guidance, funds, administrative and material support, CD conceived the project, GT, AFT, YR, CD wrote the manuscript, MA, ON, NV revised the manuscript. All authors read and approved the final manuscript.

## Acknowledgments

We thank Romain Ecalard, Lise Molimard, Myriam Sicard, Judith Hilaire and Claire Descloux for technical assistance with the animal experiments.

