## Additional file 1 for "Colon-specific immune microenvironment regulates cancer progression versus rejection"

### Additional file 1: Figure S1

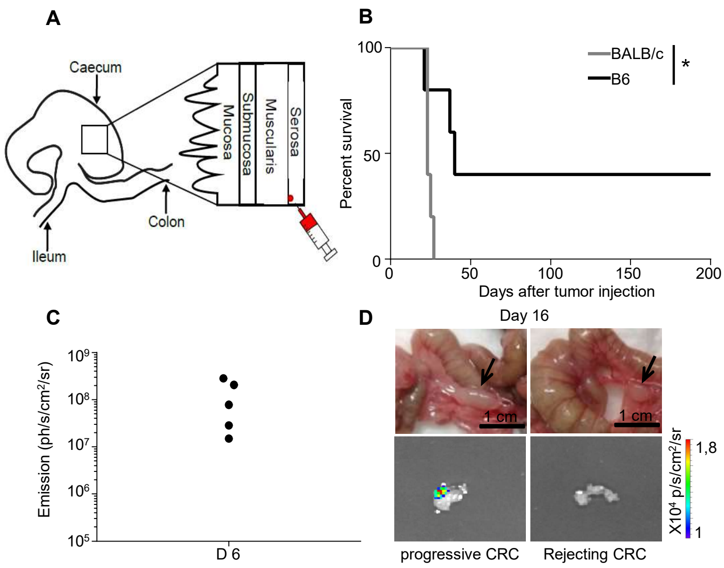

**Additional file 1: Figure S2**

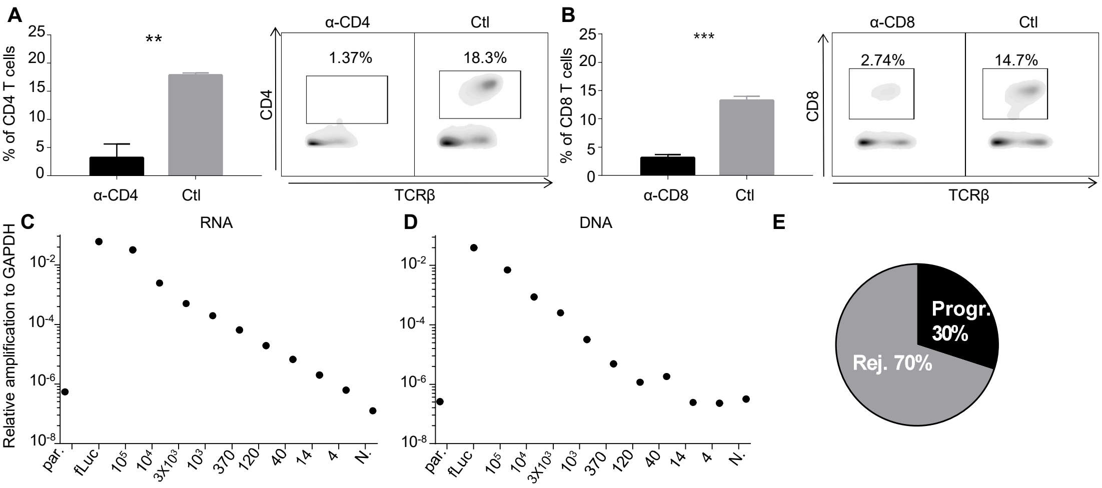

### Additional file 1: Figure S3

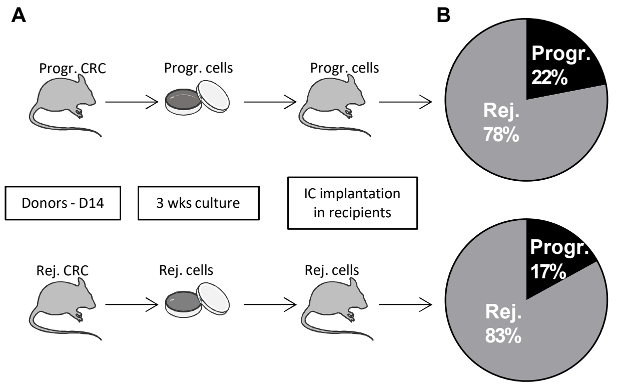

**Additional file 1: Figure S4**

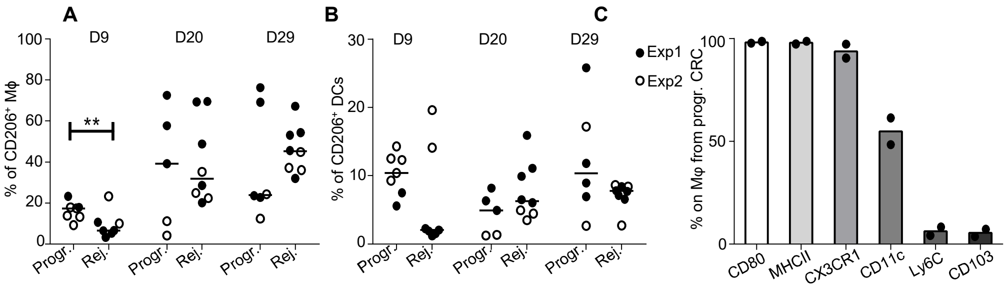

**Additional file 1: Figure S5**

A

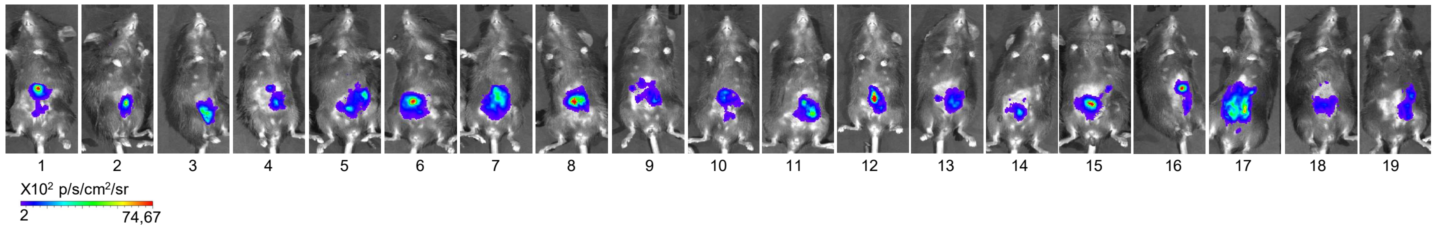


B

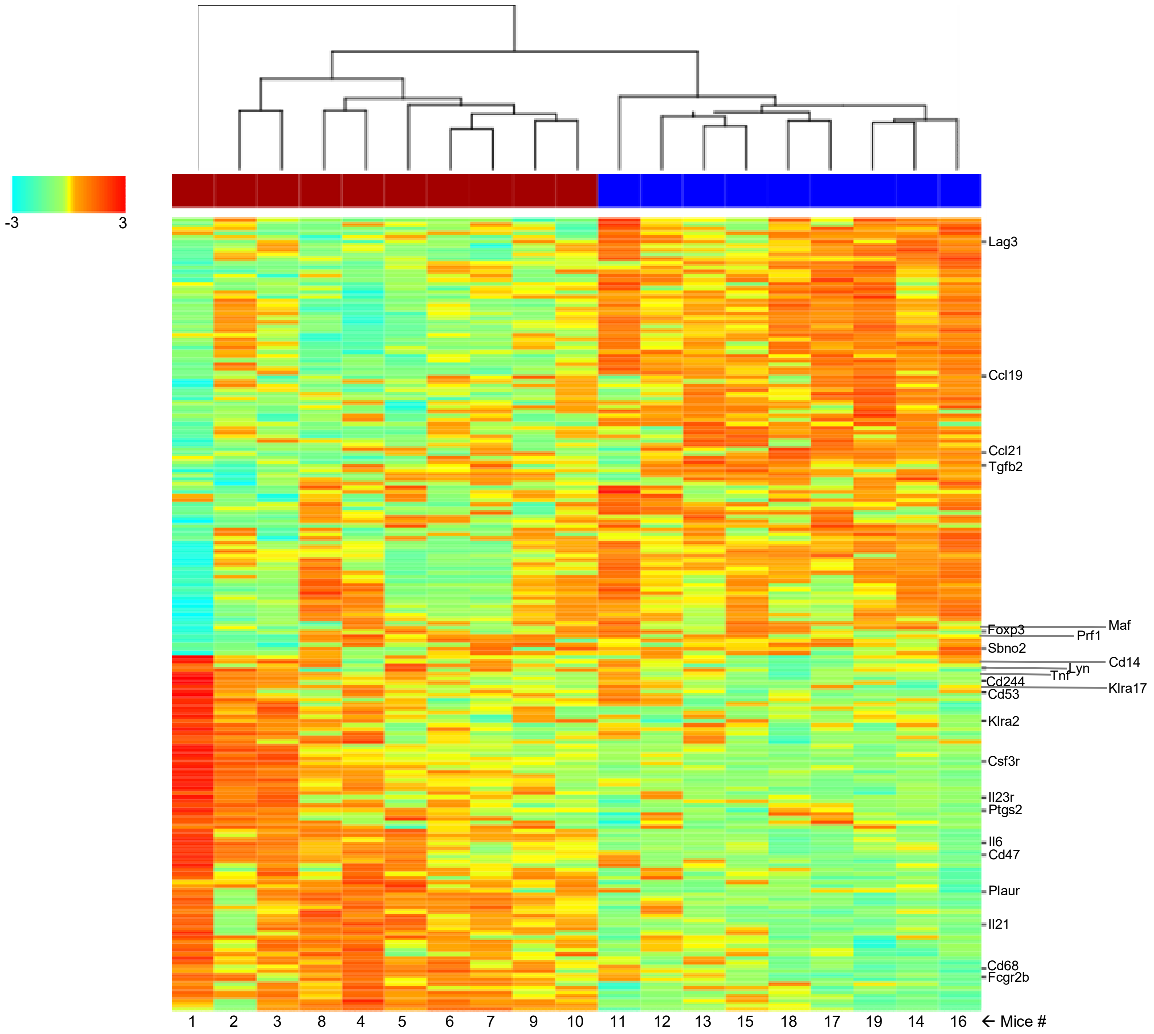
