## Additional file 2 for "Colon-specific immune microenvironment regulates cancer progression versus rejection"

**Additional file 2: Table**

| Experiment | Total number of mice | Number of proressing CRC mice | Number of rejecting CRC mice |
|---|---|---|---|
| #1 | 44 | 15 (34 %) | 29 (66 %) |
| #2 | 36 | 9 (25 %) | 27 (75 %) |
| #3 | 30 | 8 (27 %) | 22 (73 %) |
| **Total** | **110** | **32 (29 %)** | **78 (71 %)** |
