## Additional file 4 for "Colon-specific immune microenvironment regulates cancer progression versus rejection"

**Additional file 4: Supplementary Methods**

*Antibodies used in flow cytometry*

| Reactivity | Conjugate | Clone | Company | RRID Code | Dilution |
|---|---|---|---|---|---|
| CD16/CD32 | _ | 2.4G2 | BD Biosciences | RRID:AB_394655 | 1:100 |
| CD4 | PerCPCy5.5 | RM4-5 | eBioscience | RRID:AB_10717395 | 1:100 |
| CD8α | FITC | SK1 (RUO (GMP)) | BD Biosciences | RRID:AB_400279 | 1:50 |
| CD25 | PECF594 | PC61 | BD Biosciences | RRID:AB_2744346 | 1:100 |
| PD-1 | PE | J43 | BD Biosciences | RRID:AB_394284 | 1:200 |
| CD45.2 | APCCy7 | 104 (RUO) | BD Biosciences | RRID:AB_1727492 | 1:200 |
| TCRb | AF700 | H57-597 | BioLegend | RRID:AB_1027648 | 1:200 |
| FR4 | AF647 | 12A5 | BioLegend | RRID:AB_1134201 | 1:400 |
| F4-80 | PerCPCy5.5 | BM8 | eBioscience | RRID:AB_914345 | 1:100 |
| F4-80 | FITC | BM8 | BioLegend | RRID:AB_893502 | 1:200 |
| CD80 | PECF594 | 16-10A1 (RUO) | BD Biosciences | RRID:AB_2737630 | 1:200 |
| CD11c | PeCy7 | HL3 (RUO) | BD Biosciences | RRID:AB_647251 | 1:100 |
| CD11c | PE | HL3 | BD Biosciences | RRID:AB_2033996 | 1:100 |
| CD11b | PE | M1/70 | eBioscience | RRID:AB_465546 | 1:300 |
| Cd11b | BV785 | M1/70 | BioLegend | RRID:AB_2561373 | 1:200 |
| Gr1 | AF700 | RB6-8C5 | BioLegend | RRID:AB_2137487 | 1:200 |
| CD206 | BV421 | C068C2 | BioLegend | RRID:AB_2562232 | 1:100 |
| CX3CR1 | BV711 | SA011F11 | BioLegend | RRID:AB_2565939 | 1:200 |
| CD64 | PE-Dazzle | X54-5/7.1 | BioLegend | RRID:AB_2566558 | 1:100 |
| I-A/I-E | APCCy7 | M5/114.15.2 | BioLegend | RRID:AB_2069377 | 1:400 |
| CD80 | BV650 | 16-10A1 | BioLegend | RRID:AB_2686972 | 1:100 |
| CD103 | BV605 | 2E7 | BioLegend | RRID:AB_2629724 | 1:100 |
| Ly6C | eF450 | HK1.4 | eBioscience | RRID:AB_10805519 | 1:200 |

List of antibodies used for flow-cytometry

*Antibody administration for in vivo depletion*

Anti (α)-mouse CD4 and α-mouse CD8 monoclonal antibodies, kindly provided by Rémi Gence (INSERM UMR1037, UPS, Toulouse, France) were purified from rat hybridoma supernatants of GK1.5 (ATCC Cat#TIB-207, RRID:CVCL_4523) and 53-6.72 (ATCC Cat#TIB-105, RRID:CVCL_9162) cell lines, respectively. 200 µg of α-CD4 mAb or α-CD8 mAb IP administrated by 6 injections in mice (at day 61, 62, 63, 66, 68 and 73 post tumor implantation).

*RNA extraction*

Disruption of frozen tissue samples and cells was performed using a Precellys homogenizer (Bertin Instruments, Bretonneux, France). Genomic DNA or total RNA were extracted using the QIAamp DNA Mini Kit or the RNeasy Plus Mini Kit, respectively (Qiagen, Cat#51304 and 74134), according to the manufacturer's instructions.

*Luciferase qPCR*

For Reverse Transcription reaction (RT-PCR), the first cDNA strand was synthetized from 1 µg of total RNA primed with an oligo $(dT)_{15}$ Primer and reverse-transcribed with AMV Reverse Transcriptase (Reverse Transcription System - Promega, Cat# A3500). For quantitative (q)PCR, 100ng of DNA or the first cDNA strand were mixed with TaqMan Fast Advanced Master Mix (2X) and specific TaqMan qene expression assay (20X) (Mr03987587_mr, Applied Biosystems). The reactions were performed on a StepOnePlus System (Applied Biosystems) (holding stage at 50°C for 2 min and 95°C for 20 sec, followed by 45 cycles at 95°C for 1 sec, and 60°C for 20 sec). The relative expression/amplification of the gene of interest was calculated using the formula $2^{CtGAPDH-CtLuc}$ [17] and normalized to the expression/amplification level of the housekeeping gene *glyceraldehyde 3-phosphate deshydrogenase* (*GAPDH).*

*Nanostring and computational analysis*

Total RNA was isolated from day 3 fresh mouse tumors. For gene expression analysis, 100ng total RNA per reaction was used for hybridization using the nCounter PanCancer Mouse Immune Profiling panel, according to the manufacturer's nCounter XT protocol (NanoString Technologies, Panel XT_PGX_MmV1_CancerImm_CSO XT-CSO-MIP1-12 ref 115000142). The panel include 770 cancer-related mouse genes and 20 internal reference controls. Samples were processed on nCounter Sample Prep Station and nCounter Digital Analyzer (NanoString Technologies). Raw data were extracted using nSolver 2.6 software (NanoString Technologies). Gene expression values were calculated by quantile normalisation of log2-transformed data. Groups of mouse tumors were defined using principal components analysis (PCA). Differential genes expressions were tested using ANOVA and genes with pValue <0.05 (heat map

Fig 6A) or <0.005 (heat map additional file 1: Figure S5B) were selected. Z-score normalized expression of selected genes were illustrated using heat map and unsupervised clustering.
